# Connexin 30 mutation rescues high-frequency hearing, reduces gap junctional coupling and alters potassium currents in cochlear Deiters’ cells

**DOI:** 10.64898/2026.07.04.736514

**Authors:** Patrício Simões, Victoria A. Lukashkina, Andrei N. Lukashkin, Snezana Levic, Ian J. Russell

## Abstract

The early-onset, high-frequency hearing loss phenotype of CD-1 mice is rescued by the A88V mutation of the connexin 30 gap-junctional protein, despite a reduced endocochlear potential (EP), which drives cochlear hair cell receptor potentials. The mutation enables organ of Corti (OoC) extracellular receptor potentials to be similar in size to those of sensitive-hearing CBA/J mice, presumably through increased OoC resistance, despite smaller intracellular outer hair cell (OHC) receptor potentials. Low-frequency hearing in CD-1Cx30^A88V/A88V^ mice is impaired, compared with those of CBA/J and wild-type CD-1 mice. To investigate the cellular basis of OoC resistance increase and EP decrease, we made *in situ* electrophysiological measurements from Deiters’ cells (DCs) in the OoC of homozygous CD-1Cx30^A88V/A88V^ mice. DCs contribute to the OHC cytoskeletal scaffold and cochlear K^+^ recycling, and are interconnected by syncytial junctions comprising connexins 30 and 26. Measurements from CD-1Cx30^A88V/A88V^ mice were compared with those from wild-type CD-1 mice, with sensitive hearing below 12 kHz, and from the CBA/J strain. Syncytial junctional-coupling between DCs of CD-1Cx30^A88V/A88V^ mice was weaker, input resistance greater, potassium current expression was modified, and voltage-sensitive activation was shifted to more negative values compared to those of CD-1 and CBA/J mice. Inactivating potassium currents dominate in DCs of CBA/J and CD-1Cx30^A88V/A88V^ mice with excellent high-frequency hearing, and sustained currents dominate in DCs of CD-1 mice with early-onset hearing loss. These findings are discussed in relation to maintenance of OoC electrochemistry, rescue of early-onset hearing loss, impaired low-frequency hearing in CD-1Cx30^A88V/A88V^ mice, and the basis of high-frequency hearing.

**KEY POINTS SUMMARY:**

- A88V mutation in gap-junctional protein Connexin 30 reshapes cochlear electrophysiology: The mutation increases organ of Corti resistance, raises Deiters’ cell input resistance, and weakens gap-junction coupling between supporting cells. Although it reduces endocochlear potential and low-frequency hearing, it preserves extracellular outer hair cell receptor potentials, rescuing high-frequency early-onset hearing loss in wild-type CD-1 mice.
- Deiters’ cell K+-conductances in mutants resemble those of the excellent-hearing CBA/J mice: Mutant CD-1 mice show potassium-channel properties nearly identical to CBA/J strain and distinct from wild-type CD-1 mice. These faster, more sensitive K+-conductances may enhance high-frequency hearing and support resistance to hearing loss.
- Reduced gap-junction connectivity may protect against age-related hearing loss: Loss of heterotypic gap junctions and reduced supporting-cell connectivity may disrupt harmful feedback loops involved in noise-induced degeneration, explaining preserved high-frequency hearing in mutant CD-1 mice.
- Our findings provide bases for understanding mechanisms of high-frequency hearing and cochlear electrochemistry.

## INTRODUCTION

The aim of this study is to understand how the phenotypical, early onset, high-frequency hearing loss of the CD-1 mouse strain (Mahendrasingam et al., 2011; Shone et al., 1991) is rescued following the p.Ala88Val (A88V) mutation of the connexin 30 (Cx30) gap junction (GJ) protein (Bosen et al., 2014; Lukashkina et al., 2017). High-frequency hearing of the CD-1Cx30^A88V/A88V^ mouse matches that of sensitive CBA/J mice (Taberner & Liberman, 2005), although hearing below 12 kHz is impaired, even compared to that of the CD-1 background strain (Bosen et al., 2014; Kelly et al., 2019; Lukashkina et al., 2017).

Outer hair cell (OHC) receptor potentials are generated primarily by K^+^-dominated mechanoelectrical transducer (MET) current flow through the OHC MET conductance, which is modulated by sound induced displacements of the OHC hair bundles (Fettiplace, 2017) (Figure 1). MET current flow is driven by the positive endocochlear potential (EP) of the scala media (Fig. 1A) in series with the negative OHC membrane potential (Davis, 1966; Russell, 1983). Both components of this battery are smaller in CD-1Cx30^A88V/A88V^ than in CBA/J mice (Levic et al., 2022; Lukashkina et al., 2017). MET current flow across the OHC basolateral membranes, generates transmembrane potentials that control, prestin-mediated OHC electromotility (Zheng et al., 2000); Fig. 1A) which provides amplification, remarkable frequency selectivity, and sensitivity of the cochlea (Robles & Ruggero, 2001). At low sound frequencies, OHC transmembrane potentials are dominated by the intracellular receptor potential (RP). It is smaller for CD-1Cx30^A88V/A88V^ mice with reduced EP than for control mice (e.g. CBA/J) with normal EP, and accounts for their reduced sensitivity to low-frequency tones (Levic et al., 2022). Above 12 kHz, the low-pass electrical filtering of the OHC membrane electrical impedance attenuates the RP (Kössl & Russell, 1992) and OHC transmembrane voltage changes are dominated by the frequency-independent extracellular receptor potential (ERP) generated adjacent to the OHCs. ERPs are similar in size in CBA/JA and CD-1Cx30^A88V^ mice (Levic et al., 2022), despite the reduced driving voltage for the RPs and hence reduced MET current flow. Levic et al. (2022) proposed that the similar size of the ERPs in CD-1Cx30^A88V/A88V^ and CBA/J mice could be due the measured increase in the organ of Corti (OoC) resistance in CD-1Cx30^A88V/A88V^ mice. It was suggested the observed increase was due to changes in the electrophysiological properties of cochlear non-sensory cells that express Cx30 together with Cx26 in GJs. These cells include the OoC Deiters’ cells (DCs) (Jagger & Forge, 2015). DCs, and outer pillar cells, form ionically, electrically, and mechanically coupled regulatory scaffolds enclosing fluid-filled spaces surrounding the sensory-motor OHCs (Fig. 1A, e.g. (Lukashkina et al., 2024; Zhou et al., 2022). We made *ex vivo* electrophysiological measurements from DCs to see if the increased resistance of the OoC, whose GJ plaque sizes are reduced in CD-1Cx30^A88V/A88V^ mice (Kelly et al., 2019), was associated with electrophysiological changes in the GJs (Levic et al., 2022).

**Figure 1.**
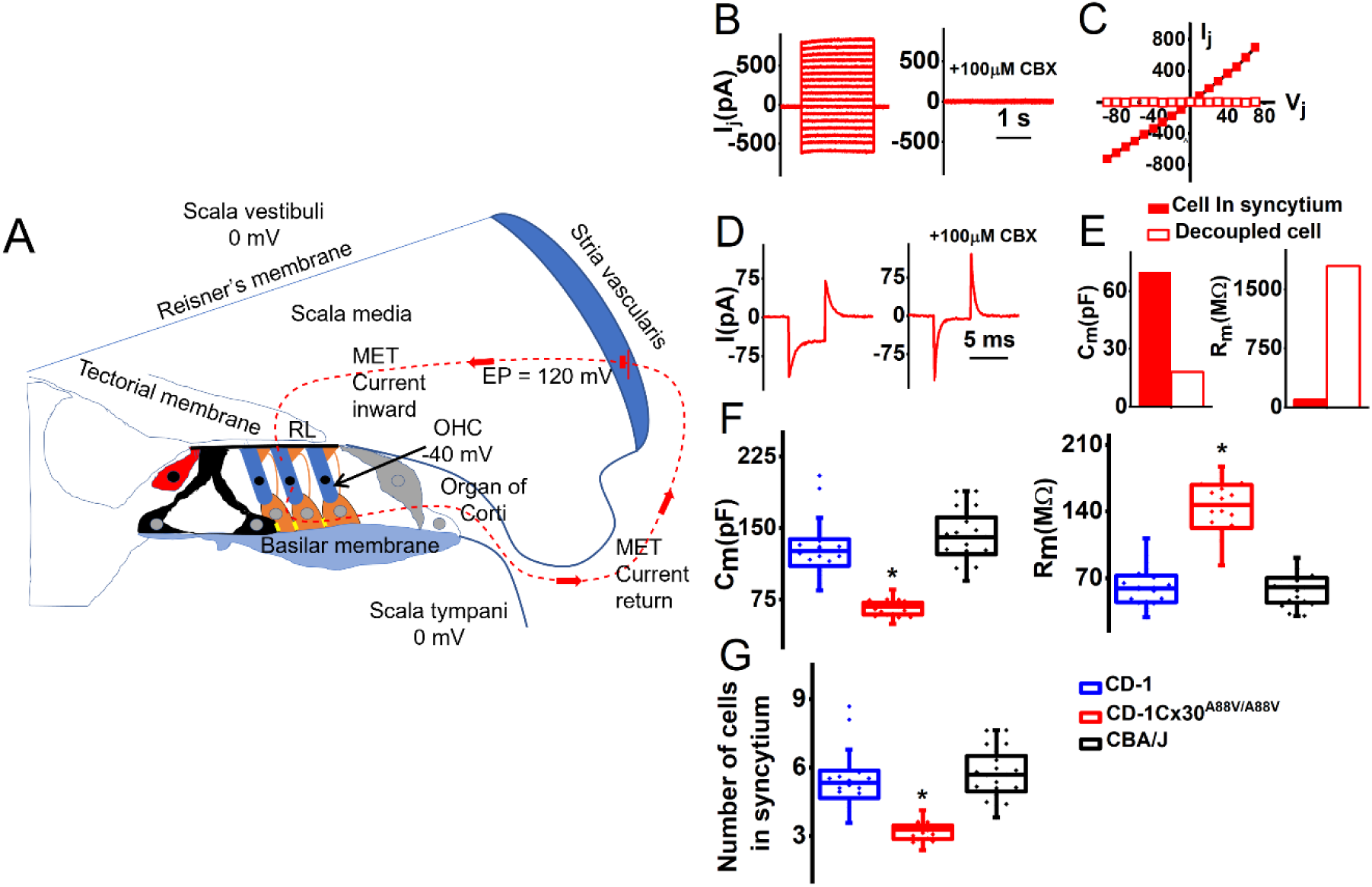
Addition of carbenoxolone decouples Deiters’ cells from the organ of Corti syncytium electrical network and reveals that the electrical properties of gap junctions in CD-1 and CBA/J Deiters’ cells are different from those of CD-1Cx30^A88V/A88V^ mice. **A**. Schematic cross-section of the cochlea and outer hair cell (OHC) mechanoelectrical transduction (MET) current pathway. OHCs (dark blue), Deiters’ cells (orange) with phalangeal processes extending to OHC apical surface, and the reticular lamina (RL). Inner hair cells (red), pillar cells (black). Schematic of inward and return pathways of the MET current (continuous red dashed line) driven by the in-series batteries of the positive endocochlear potential (EP) and negative OHC resting membrane potential. **B**. An example of junctional currents (I_j_) recorded from a single Deiters’ cell (DC) in the syncytium in CD-1Cx30^A88V/A88V^ mice in response to voltage steps delivered in 10 mV increments, from holding potential of -40 mV between -100mV to +70 mV before (left) and after addition of 100 μM CBX (right). **C.** Plot of junctional currents (I_j_) in response to junctional potential changes (V_j_) measured at the peak current before (filled symbols) and after addition of 100 μM CBX (open symbols) for the same cell shown in B. **D.** An example of a capacitive transient recorded from a single DC cell in response to the - 5 mV step from the holding potential of -40 mV before (left) and after addition of 100 μM carbenoxolone (CBX), a gap junction blocker (right). The same cell as in B. **E**. Plot of the membrane capacitance (C_m_) and membrane resistance (R_m_) for the same cell as in B before (solid bars) and after addition of 100 μM CBX (open bars). **F**. Left. Box plot C_m_ (pF, mean ± SD, number of measurements, n) for cells within the syncytium measured in CD-1 (129.5 ± 30.8, n=20), CD-1Cx30^A88V/A88V^ (66.5 ± 8.5, n=20), and CBA/J (140.9 ± 27.1, n=20) mice. Right. Box plot of R_m_ (MΩ, mean ± SD, number of measurements, n) for cells within the syncytium measured in CD-1 (60.6 ± 20.9, n=20), CBA/J (58.2 ± 16.8, n=20), and CD-1Cx30^A88V/A88V^ (144.8 ± 27.9, n=20) mice. **G**. Box plot of estimated number of coupled cells (mean ± SD, number of measurements, n) in CD-1 (5.5 ± 1.3, n=20), CBA/J (5.7 ± 1.1, n=20) and CD-1Cx30^A88V/A88V^ (3.2 ±0.4, n=20). The number of coupled DCs was calculated dividing C_m_ measured for cells within the syncytium (Fig.1F, left) by the mean single-cell capacitance C_m_ measured in the presence of 100 μM CBX.

DCs are attributed with regulating Cortilymph electrochemistry (Jagger & Forge, 2015) including K^+^ uptake and recycling. Cortilymph K^+^ levels increase during sound stimulation (Johnstone et al., 1989) and K^+^ recycling is essential for maintaining EP and cochlear fluid homeostasis (Kim et al., 2021; Köppl et al., 2018). Zdebik et al. (2009) proposed that K^+^ exits from OHCs via KCNQ4 K^+^ channels and accumulates in Cortilymph between the OHCs and DCs. K^+^ is taken into the DCs by KCC4 and KCC3 (K^+^-Cl^−^cotransporters) and K^+^ moves through GJs to adjacent epithelial cells and into the scala media. Outward rectifying K^+^ currents recorded from guinea pig DCs (Chung et al., 2013; Nenov et al., 1998) have been postulated to be important in repolarizing DCs during K^+^ buffering of the Cortilymph and K^+^ recycling (Yang & Wang, 2002). Therefore, another aim of this study is to characterize differences in DC K^+^ currents in *ex vivo* preparations of different mouse strains and to understand their potential roles in Cortilymph K^+^ homeostasis, and how differences might account for early-onset-high-frequency-hearing-loss and its rescue by the Cx30 A88V mutation.

## MATERIALS AND METHODS

### ETHICAL APPROVAL

All in vitro and in vivo experiments performed at the University of Brighton complied with Home Office guidelines under the Animals (Scientific Procedures) Act of 1986 and were approved by the University of Brighton Animal Welfare and Ethical Review Body (PP3408916). All of this research adhered to the ethics principles and policies of *The Journal of Physiology*. Every effort was made to reduce the number of animals used and minimize animal suffering. Approximately equal numbers of male and female mice were used for each set of experiments.

Male and female mice were housed in the University of Brighton Bioresources Unit with a 12 h light/dark cycle and a free access to food and water. All mice used here were originally purchased from the Jackson Laboratory. Pups were housed with the dams and allowed to nurse freely until used for experiments.

### TISSUE PREPARATION

The pups were decapitated (Schedule 1). The cochlea was freshly isolated and dissected using an ice-cold solution containing in mM: NaCl 135, KCl 5.8, CaCl_2_ 1.3, MgCl_2_ 0.9, NaH_2_PO_4_ 0.7, d-glucose 5.6, HEPES 10, Sodium pyruvate 2, pH 7.5 (adjusted with 1M NaOH), osmolarity ∼ 308 mOsm. DCs were studied in acutely dissected OoC from postnatal day 14 (P14) to P20, where the day of birth is P0. The dissected apical coil of the OoC was transferred to a microscope chamber, immobilized using a nylon mesh fixed to a stainless-steel ring and viewed using an upright microscope (Axioskope, Zeiss, Germany). Cells in the dissected tissue were observed with Nomarski differential interface contrast optics (40× water immersion objectives).

### ELECTROPHYSIOLOGY

All currents were recorded in a whole-cell voltage-clamp configuration. The patch pipettes were fabricated (PC-10, Narishige, Japan) using borosilicate glass (Sutter Instruments), with 1.5 mm O.D., and heat polished (MF-830, Narishige, Japan). The pipette resistance in the extracellular solution was 2-5 MΩ. Single DCs or DC pairs were voltage clamped using a dual Axopatch 700B amplifier (Molecular Devices, Union City, CA), and the response was filtered at a frequency of 2–5 kHz through a low-pass Bessel filter. The data were digitized at 5–500 kHz using an analog-to-digital converter (Digidata 1500; Molecular Devices). The sampling frequency was determined by the protocol used. The liquid junction potentials were measured (3.5±0.9 mV, n= 149) and corrected online (Neher, 1992). Current recordings were collected using pCLAMP software (version 10, Axon Instruments, Foster City, CA, USA). Data were analysed using pClamp10 (Molecular Devices), Origin9.1 (OriginLab Corp. Northampton, MA) and Excel (Microsoft). Significant differences between groups were tested using ANOVA and post hoc Tukey test, with *p* < 0.001, indicating a statistically significant difference.

### Gap junctional recordings

All recordings were done at room temperature using extracellular solution containing (in mM) NaCl 100, TEA-Cl 20, CsCl 20, BaCl_2_ 1.25, MgCl_2_ 1.48, HEPES 10, pH 7.2 and osmolarity ∼ 300 mOsm. Patch pipettes were filled with the intracellular solution containing in mM: CsCl 140, EGTA 5, MgCl_2_ 2, HEPES 10, pH 7.2, osmolarity ∼ 290 mOsm. All chemicals were obtained from Sigma Aldrich, UK.

#### Estimating the number of cells coupled electrically in the syncytium by measuring cell capacitance and membrane resistance

We measured capacitive currents to estimate the membrane capacitance (C_m_) from a holding potential of -40 mV, using 5 mV hyperpolarizing steps (5 ms duration) when no ionic currents were activated. These measurements were made from cells electrically coupled to the syncytium and from cells that were effectively electrically isolated from the syncytium using 100 μM carbenoxolone (CBX), which has been shown to block gap junctional currents in the inner ear (Abitbol et al., 2020; Schütz et al., 2010).

C_m_ was calculated by dividing the area under the transient current in response to a voltage step as described (Levic et al., 2007). The capacitive decay was fitted with a single exponential curve to determine the membrane time constant. Uncompensated series resistance (R_s_) was estimated from the membrane time constant, given its capacitance. This study included ∼120 cells with R_s_ within a 5 – 10 MΩ range. After 60–90% compensation of the mean residual, uncompensated R_s_ was 4.1 ± 0.6 MΩ. The seal resistance was typically 1 – 5 GΩ. Whole-cell membrane resistance (R_m_) was calculated from measured current amplitude and voltage step magnitude (5 mV) by dividing the voltage step by current amplitude.

#### Syncytial junctional current recordings

To qualitatively assess the electrical coupling between DCs in the syncytium, we used the double voltage-clamp technique to measure junctional conductance (G_j_) between adjacent cells in the syncytium (Ma et al., 2016; Meme et al., 2009). Our objective was to discover if the syncytial junction conductance of DCs was different between the CD-1, CBA/J and CD-1Cx30^A88V/A88V^ mouse strains. The outcomes of our measurements pointed in the direction of the basis for differences between strains. All conductances and capacitances within the syncytium contribute uncontrollably to the recorded currents. Accordingly our values likely underestimate the true syncytial input resistance, capacitance, and individual junctional conductances (Stephan et al., 2021). While this approach differs from measurements made from single or pairs of isolated cells, it successfully highlights qualitative differences in the electrical coupling between DC syncytia across different mouse strains, an approach previously validated in studying astrocyte gap junctional coupling *in situ* (e.g. (Meme et al., 2009; Xu et al., 2010), reviewed in (Stephan et al., 2021). Any observed differences could be due to the number of functionally connected cells, the specific connexin channels forming the GJs and the biophysical properties of the DCs. The exact bases of the differences we have discovered are the subject of further investigations that are beyond the aims of the current study. The measurements, however, provide us with a qualitative description of strain-dependent syncytial connectivity because the measurements in the different strains were done under the same experimental conditions. Thus, two neighbouring cells were held at -40 mV. Test voltages ranging from -110 mV to +60 mV were applied to one cell (cell 1) and the other cell (cell 2) was held at a constant voltage of -40 mV. Estimates of the junctional potential between cells in the syncytium (V_j_) was calculated as

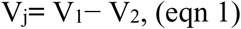

where V_1_ is the applied voltage in stepped cell 1 (from -110 to +60 mV), and V_2_ is the holding voltage in non-stepped cell 2 (−40 mV).

Syncytial junctional currents (I_j_) were measured in a cell 2, as

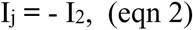

where I_2_ is current response recorded in cell 2.

For each cell pair within a syncytium, the syncytial junctional conductances measured from a particular cell (instantaneous, G_j.inst_ and steady state, G_j.ss_) were calculated as

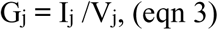

where I_j.inst_ was measured ∼ 5ms after the beginning of the step pulse, and I_j.ss_ was measured at ∼1s at the end of voltage step (Spray et al., 1981).

### Potassium current recordings

All recordings were done at room temperature. Potassium currents were recorded using the same solution as for the dissection of single cells electrically isolated from the syncytium using CBX. Patch pipettes were filled with intracellular solution containing in mM: KCl 131, MgCl_2_ 3, Na_2_ATP 5, Na_2_-phosphocreatine 10, EGTA 1, HEPES 10, Na_2_GTP, pH 7.3 (adjusted with 1M KOH), osmolarity ∼ 293 mOsm. All chemicals were obtained from Sigma Aldrich, UK.

Cell membrane capacitance C_m_, membrane resistance R_m_ and series resistance R_s_ were calculated as described above. This study included 149 cells with R_s_ within a 5–15 MΩ range. After 60–90% compensation of the mean residual, uncompensated R_s_ was 5.1±0.5 MΩ. The seal resistance was typically 2-5 GΩ. No online leak current subtraction was made, and only recordings with holding currents less than 50 pA were accepted for analyses. The number of cells (n) is given for each data set. Time constants for potassium current activation/deactivation (τ_i_) were obtained from fits using Origin9.1 software (OriginLab Corp. Northampton, MA). Time constants were obtained by fitting multiple exponential terms to the activation and decay of the current. The fitting equation was

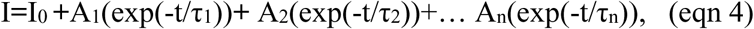

where I_0_ is the initial current magnitude, τ_1_, τ_2_…τ_n_ are the time constants, and A_1_, A_2_…A_n_, are the proportionality constants. The steady-state inactivation curve was generated from normalized currents measured at a test potential following several conditioning pre-pulses. Pooled data were presented as mean ± SD. Statistical comparisons were made using two-tailed Student’s *t-*test. In multiple comparisons tests, significance was adjusted with the Dunn–Sidak correction, where α’=1-(1-α)^1/k^, where k is the number of comparisons performed between the same groups (Zar, 1999).

## RESULTS

### Electrical syncytial coupling between Deiters’ cells is reduced in CD-1Cx30^A88V/A88V^ compared with CBA/J and CD-1 mice

DCs form a syncytium in the OoC and, as a first approximation to investigating the properties of syncytial coupling in DCs, we measured the cell capacitance (C_m_) and resistance (R_m_) of cells within the syncytium and cells electrically isolated from the syncytium using 100 μM carbenoxolone (CBX). CBX has been shown to effectively block GJ currents in the inner ear (Abitbol et al., 2020; Schütz et al., 2010).

We were able to confirm that the currents recorded from a single cell, as shown in Fig. 1B,C while simultaneously delivering a voltage step to a neighbouring cell, are due to junctional conductances because they were also sensitive to CBX block in the presence of pharmacological blockers of other known voltage gated conductances found in DCs (Methods). This approach enabled us to estimate the contribution of syncytial coupling to the capacitance and resistance measurements by comparing them to measurements from functionally isolated single cell using CBX block as illustrated in Fig. 1D-F.

CBX application decreased C_m_ (Fig. 1D-E) with simultaneous increase of R_m_ (Fig. 1E). The mean calculated C_m_ for DCs in the syncytium shown in Fig. 1F (pF, mean ± SD, number of measurements, n) was (129.5 ± 30.8, n=20) for CD-1, (140.9 ± 27.1, n=20) for CBA/J, and (66.5 ± 8.5, n=20) for CD-1Cx30^A88V/A88V^. The post-hoc Tukey test reveals a significant difference in syncytial C_m_ between CD-1 and CD-1Cx30^A88V/A88V^ (p < 0.0001), and between CBA/J and CD-1Cx30^A88V/A88V^ (p < 0.0001), but not between CD-1 and CBA/J (p = 0.3), indicating reduced electrical coupling between DCs in CD-1Cx30^A88V/A88V^ mutants (Santos-Sacchi, 1991). By blocking the GJ conductances in the syncytium with CBX, individual cells were electrically isolated (Fig. 1B, C), thereby providing an opportunity to estimate their single-cell C_m_. The mean DC single-cell C_m_ (pF, mean ± SD, number of measurements, n) was (23.6 ± 11.6, n = 46) for CD-1, (24.7 ± 11.8, n = 50) for CBA/J, and (20.7 ± 9.2, n = 53) for CD-1Cx30^A88V/A88V^. No significant differences were observed in single-cell C_m_ of DCs between the three mouse strains (analysis of variance (ANOVA, p = 0.417), suggesting that the cell sizes were similar in different strains.

We also used the measurements of C_m_ in a syncytium (Fig. 1F) to make a rough estimate the number of functionally connected cells (Fig. 1G), given the known single-cell C_m_. The number of coupled DCs is calculated by dividing the mean syncytial C_m_ by the mean single-cell C_m_ measured in the presence of 100 μM CBX. Our estimate is likely to be an underestimate of the number of coupled cells. This is because the strength of electrical coupling between cells dissipates over distance (e.g. (Pang et al., 2024)). Thus, based on our approach, some of the estimated coupled cells may include coupled distal cells that make only small contributions to the measured overall syncytium capacitance. Nevertheless, we applied the same method, assuming equal and full capacitance contribution of all cells when comparing the three strains. The mean calculated number of electrically connected cells within a syncytium shown in Fig. 1G (mean ± SD, number of measurements, n) was (5.5 ± 1.3, n=20) for CD-1, (5.7 ± 1.1, n=20) for CBA/J, and (3.2 ±0.4, n=20) for CD-1Cx30^A88V/A88V^. Post-hoc Tukey test reveals significant differences between CD-1 and CD-1Cx30^A88V/A88V^(p < 0.0001) and CBA/J and -1Cx30^A88V/A88V^ (p < 0.0001), but not between CD-1 and CBA/J (p =0.77). Thus, our data indicates that either fewer cells are coupled in CD-1Cx30^A88V/A88V^ mice compared with the other two strains, or that the interconnections are weaker.

The mean calculated R_m_ within a syncytium are shown in Fig. 1F. They (MΩ, mean ± SD, number of measurements, n) were (60.6± 20.9, n=20) for CD-1, (58.2 ± 16.8, n=20) for CBA/J, and (144.8 ± 27.9, n=20) for CD-1Cx30^A88V/A88V^. The post-hoc Tukey test reveals significant differences between CD-1 and CD-1Cx30^A88V/A88V^ (p < 0.0001) and CBA/J and CD-1Cx30^A88V/A88V^ (p = 0.0001), but not between CD-1 and CBA/J (p = 0.94). Our results indicate that the strength of electrical coupling between DCs cells within a syncytium is significantly reduced while the input resistance R_m_ is increased in CD-1Cx30^A88V/A88V^ mutants.

Together, this data shows that the functional coupling within DCs in CBA/J and CD-1 mice is limited to about 5-6 cells suggesting that the supporting cells are probably not acting like an isopotential syncytium (Lagostena et al., 2001). Moreover, this coupling seems to be further reduced due to the Cx30^AV88/AV88^ mutation (Fig. 1G).

### Heterogeneous, syncytial junctional, voltage coupling between Deiters’ cells was observed in CD-1 and CBA/J but not in CD-1Cx30^A88V/A88V^mice

We further investigated the effects of the Cx30^A88V/A88V^ mutation on electrical coupling between adjacent pairs of DCs within the syncytium of the isolated OoC. The electrical network of the syncytium more closely resembles the *in vivo* situation than that of pairs of isolated DCs (Santos-Sacchi, 2000; Stephan et al., 2021). The unknown extent of electrical interconnectivity between cells comprising the syncytium and the potential sensitivity of the electrical network to the ionic and electrical extracellular environment (Santos-Sacchi, 1987, 2000) makes interpretation of the outcomes more complex and difficult than that of the electrical coupling between two isolated DCs. However, isolated OoCs from the different mouse strains were subjected to identical recording conditions which makes it possible, using this qualitative approach, to address issues raised by the outcomes of the *in vivo* measurements (Levic et al., 2022).

Previous studies demonstrated that, in addition to the homotypic GJ conductance (HomC) with symmetric I-V characteristic, most GJ conductance in the OoC have asymmetric I-V characteristics, which has been taken to indicate heterogeneous coupling, perhaps resulting from heterotypic GJ conductance (HetC) or, possibly, heteromeric configurations (Zhao, 2000; Zhao & Santos-Sacchi, 2000). Pairs of adjacent DCs were initially clamped to -40 mV, which was chosen as the most stable potential producing minimal leak currents. Syncytial junctional voltage (V_j_, calculated as detailed in the Methods) was produced by 2-second voltage steps from -110 mV to +60 mV applied to cell 1 (V_1_, 10 mV increments), and cell 2 was continuously held at -40 mV (V_2_). The change in V_j_ = V_2_ – V_1_ was thus from -100 to +70 mV. Typical examples of syncytial junctional currents (I_j_) recorded during 2-second pulse stimulation from pairs of CD-1, CD-1Cx30^A88V/A88V^ and CBA/J DCs are shown in Fig. 2A.

**Figure 2.**
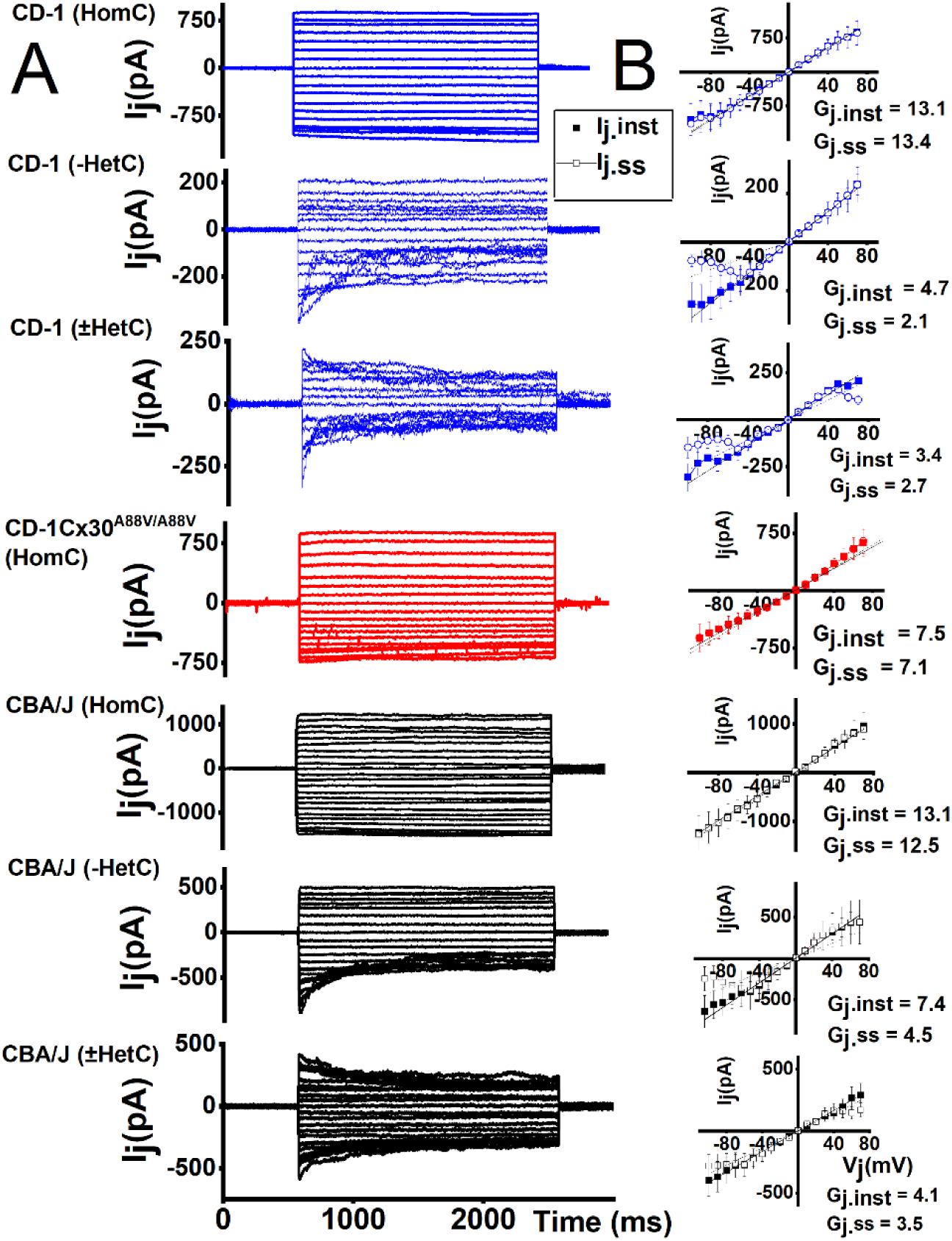
Heterogeneous transjunctional voltage coupling was observed in CD-1 and CBA/J but not in CD-1Cx30^A88V/A88V^ Deiters’ cells. **(A)** Typical examples of transjunctional currents (I_j_) recorded during 2-s pulse stimulation from -100 to +70 mV from pairs of CD-1, CD-1Cx30^A88V/A88V^ and CBA/J Deiters’ cells (DCs). Cells were initial held at -40 mV and step voltage was applied to one cell and the transjunctional currents were measured in the non-stepped adjacent cell. CD-1 and CBA/J cells, but not CD-1Cx30^A88V/A88V^, showed a heterogenous transjunctional voltage (V_j_) and time dependence, defined by three different categories of response (HomC, -HetC, ±HetC). **(B)** Summary plots of I_j_ in response to V_j_ stimulation obtained from the DC pairs, as classified above. Instantaneous (I_j.inst_) and steady-state (I_j.ss_) currents were measured at onset (5 ms) and end (∼1sec) of voltage steps and are represented by solid squares and open circles, respectively. Values of instantaneous and steady-state gap junctional conductances (G_j_) correspond to the slope of the linear regression of the represented I_j_-V_j_ relationships. Please note differences in the vertical axes that enable comparisons to be made between the time-different dependencies of the I_j_.

Syncytial junctional currents (I_j_) in the DCs of CD-1Cx30^A88V/A88V^ mice are homogenous with little time-dependence (Fig. 2A, red traces). In contrast, I_j_ in CD-1 and CBA/J (Fig. 2A, blue and black traces) DCs revealed heterogenous voltage coupling and time dependence. We were able to separate the DC responses of CD-1 mice into 3 categories based on the voltage and time dependence of I_j_. CD-1(HomC) I_j_ displayed no or little voltage-and time-dependent inactivation. CD-1(−HetC) I_j_ showed pronounced voltage- and time-dependent inactivation when the V_j_ is lower than -50 mV. CD-1(±HetC) I_j_ displayed pronounced voltage- and time-dependent inactivation when V_j_ is more negative than -50 mV and more positive than 50 mV (Fig. 2A). These two last categories also exhibited lower I_j_ magnitudes than CD-1 and CBA/J (HomC). Fig. 2B shows the summary plots of I_j_ in response to V_j_ stimulation obtained from syncytially coupled DC pairs using the above classification. The peak instantaneous (I_j.inst_) and steady-state (I_j.ss_) transjunctional currents were measured at the onset (∼5 ms) and the end of each voltage step (∼2 sec), respectively. In all cell types, the I_j.inst_ relative to the V_j_ was approximately linear; I_j.ss_ showed, however, heterogenous deviation from linearity in CD-1 and CBA/J cells for potentials more negative than -50 mV (in CD-1 and CBA/J(−HetC)) and more positive than 50 mV (in CD-1 and CBA/J(±HetC)).

The comparison of the instantaneous syncytial conductances of DCs across different strains (Fig. 3) shows that the conductance of non-inactivating I_j_ (HomC) in CD-1Cx30^A88V/A88V^ is 40-50% of that of CD-1(HomC). Measured at +70 mV, it was (mean ± SD, n = 6) 8.9 ± 0.7 nS for CD-1Cx30^A88V/A88V^(HomC) and 12.5 ± 1.2 nS for CD-1(HomC). CD-1Cx30^A88V/A88V^(HomC) conductance is, however, 110-160% higher than the conductance for inactivating I_j_ (CD-1(−HetC) and (±HetC)). Measured at +70 mV, the conductance was (mean ± SD, n = 6) 5.1 ± 0.6 nS for CD-1(−HetC) and 2.9 ± 0.3 nS CD-1(±HetC). At the DC resting membrane potential (∼110 mV), the CBA/J(HomC) conductance, which is observed in about 75% of the DC pairs, is twice as large as the CD-1Cx30^A88V/A88V^(HomC) syncytial gap junctional conductance.

**Figure 3.**
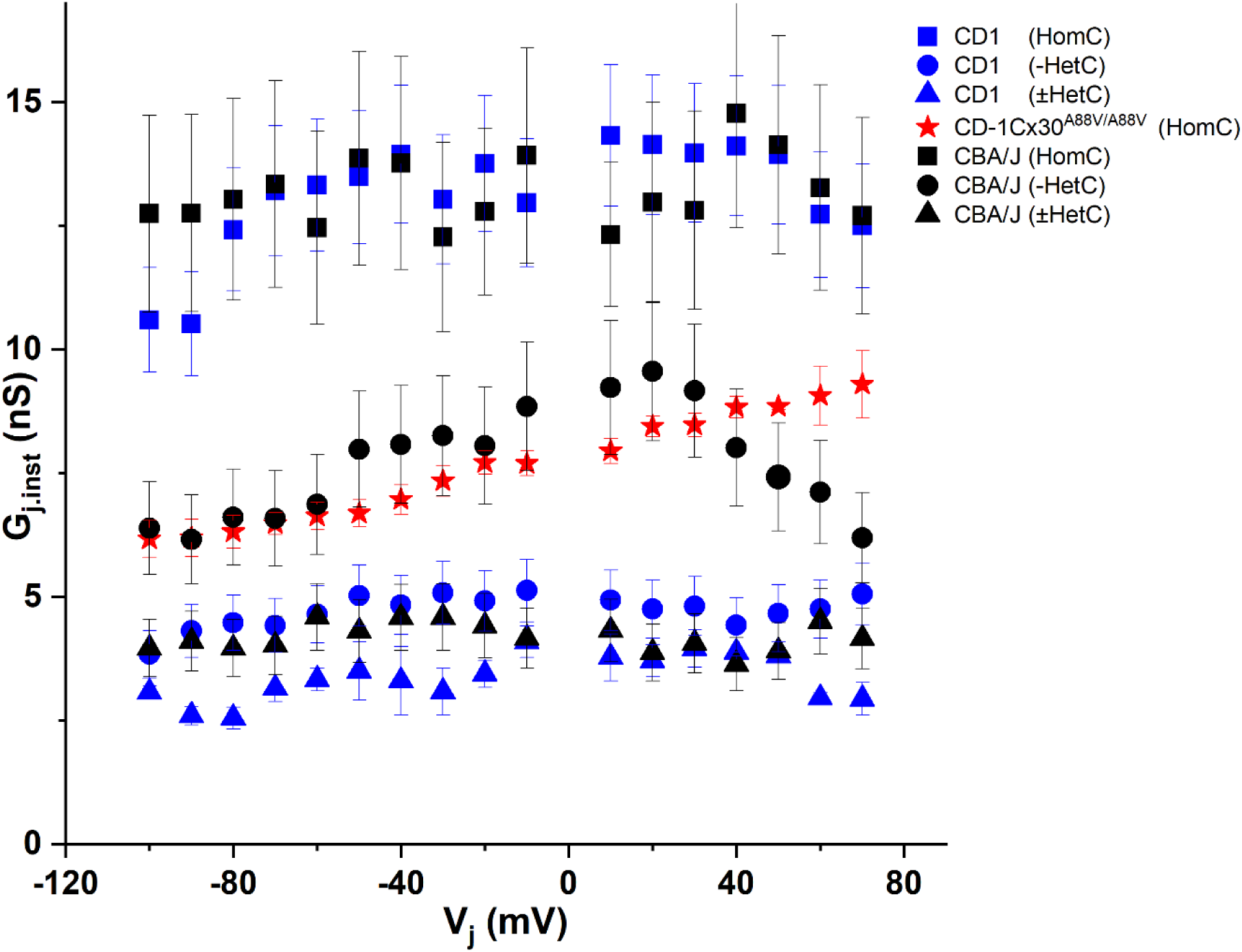
Summary comparison of the Deiters’ cells gap junctional conductance values in CD-1, CBA/J and CD-1Cx30^A88V/A88V^ cells, as determined from I_j_ vs V_j_ instantaneous plots, reveal voltage independent coupling (HomC) is similar in CD-1 and CBA/J cells, and significantly higher than that in CD-1Cx30^A88V/A88V^ cells. The conductance of voltage independent coupling is significantly greater than that of voltage dependent (Het) coupling. However, the conductance of -Het coupling in CBA/J cells is greater than in CD-1 cells.

### Vj-dependent gating in Deiters’ cell syncytium shows dependence on direction of stimulation in coupled cells, producing symmetrical responses in CD-1Cx30^A88v/A88V^ and asymmetrical responses in CD-1 and CBA/J strains

The syncytial junctional currents obtained from patching two adjacent DCs were investigated by first stimulating one cell in a pair with voltage steps and recording the I_j_ in the second cell, and then stimulating the second, while recording in the first cell of the pair (Fig. 4, top panels). The response of CD-1Cx30^A88V/A88V^ DC pairs to transjunctional reciprocal stimulation typically produced homogenous and symmetrical I_j,_ that showed minimal time dependence (Fig. 4, red). The response of CD-1 and CBA/J DC pairs to transjunctional reciprocal stimulation was heterogenous, with some pairs producing strong, symmetrical I_j_, whereas other pairs produced asymmetrical responses with V_j_-dependent rectifying currents (Fig. 4, blue and black respectively). These results suggest that the connexin channel composition is heterotypic in the syncytia of CD-1 and CBA/J DCs, whereas the syncytia of CD-1Cx30^A88V/A88V^ DCs is formed through homotypic GJs.

**Figure 4.**
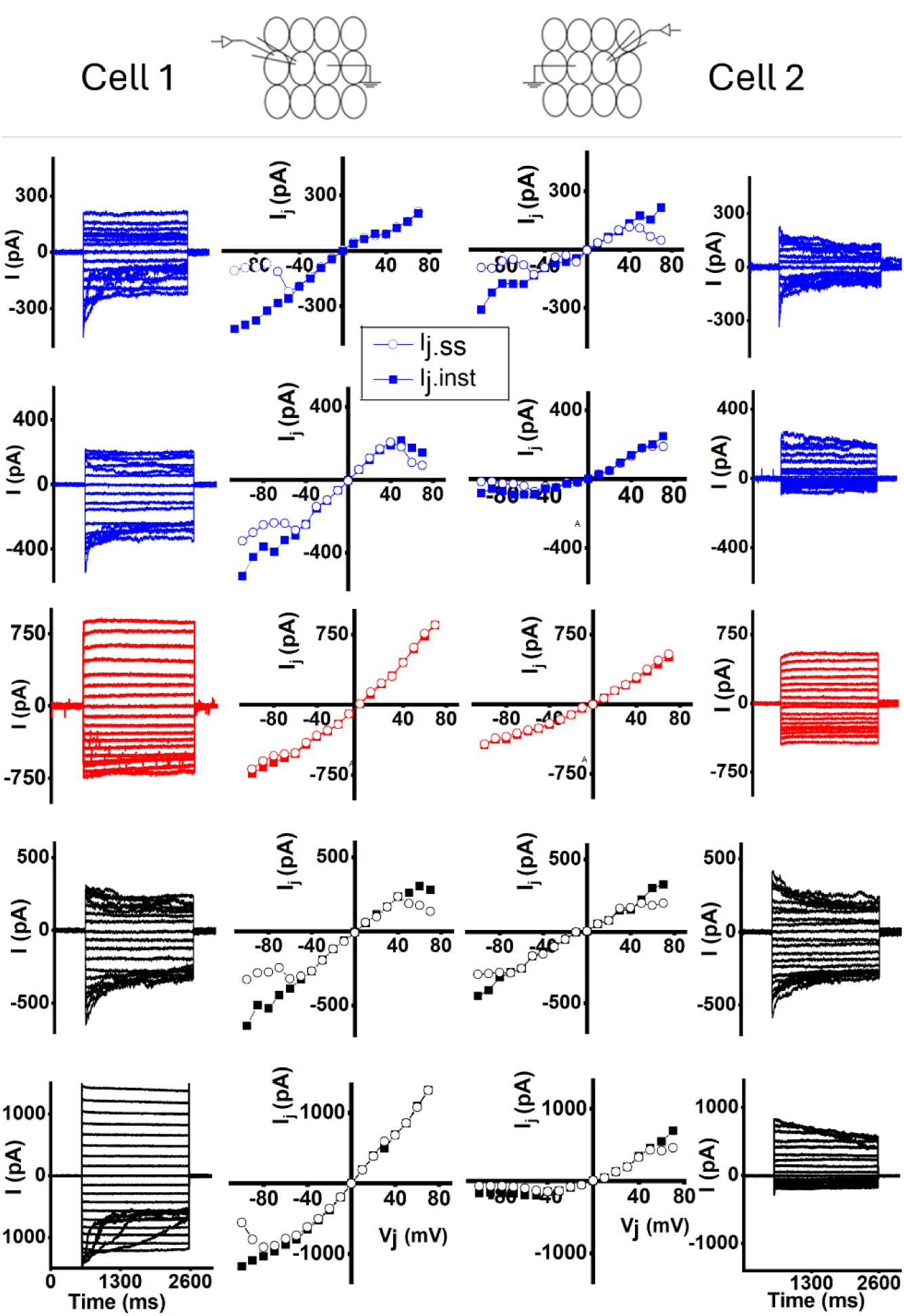
Vj-dependent gating in the same syncytial cell pairs shows dependence on direction of stimulation in coupled cells, producing asymmetrical responses in CD-1 (blue) and CBA/J (black) Deiters’ cells and symmetrical responses in CD-1Cx30^A88V/A88V^ (red) cells. Examples of syncytial responses elicited in different coupled Deiters’ cell pairs. (Cell 1) Plots of I_j_ response in cell 1 to voltage stimulation in cell 2, and (Cell 2) I_j_ response in cell 2 to voltage stimulation in cell 1. Instantaneous (I_j.inst_, solid squares) and steady-state (I_j.ss_, open circles) were measured at the onset (∼5 ms) and the end of each voltage step (∼2 sec), respectively.

### Kinetics of inactivation of potassium currents measured in situ in Deiters’ cells of CD-1Cx30^A88V/A88V^ mice resemble the kinetics in CBA/J mice with good high-frequency hearing

To further investigate the cellular basis for differences in the *in vivo* electrical responses that were measured in supporting cells of the OoC of CBA/J and CD-1Cx30^A88V/A88V^ mice (Levic et al., 2022), potassium currents were measured in DCs from *in situ* isolated middle-turns of the cochlea of CBA/J, CD-1Cx30^A88V/A88V^ and CD-1 mice. CBA/J mice, with excellent high-frequency hearing, were used as the “WT control” in the *in vivo* measurements because CD-1 mice express rapid, early onset hearing loss (Shone et al., 1991). Measurements were also made from DCs of CD-1 mice to discover if, and how, the Cx30^A88V/A88V^ mutation altered the electrophysiological properties of the DCs in the CD-1 background strain, and if these changes provided insight to changes in potassium currents that might be associated with the preservation of high-frequency hearing. The ages of the mice were restricted to ∼ P20 to avoid the effects of rapid-onset-hearing loss that appear after this time in CD-1 mice.

Potassium currents were elicited in DCs using 500 ms voltage steps in 10-mV increments from -90 to +50 mV, from a holding potential of -80 mV. Representative current traces from CD-1, CD-1Cx30^A88V/A88V^ and CBA/J mice are shown in Fig. 5 A, B and C, respectively. DC potassium currents were significantly different in amplitude, density, and voltage dependence of activation between CD-1, CBA/J, and CD-1Cx30^A88V/A88V^ mice (Figs. 5 D, E, F, respectively). At an activation voltage of +50 mV, the average DC potassium current amplitude of CD-1 mice (5430 ± 724 pA) was significantly larger than either those of CBA/J (3513 ± 468 pA) or CD-1Cx30^A88V/A88V^ (2829 ± 416 pA) mice (Fig. 5 D, t = 11.11, P < 0.0001, d.f.= 48, α’=0.017). These differences were also reflected in the current density (Fig. 5E). Current density at +50 mV depolarization of both CD-1Cx30^A88V/A88V^ (334 ± 45 pA/pF) and CBA/J (270 ± 40 pA/pF) was significantly lower than in CD-1 mice (517±68 pA/pF; t = 11.16, P < 0.0001, d.f.= 48, α’=0.017; Fig. 5E). Differences in the current density were not due to differences in cell size because cell capacitance was similar in CD-1, CD-1Cx30^A88V/A88V^, and CBA/J being 11.3 ± 0.4 pF, 11.6 ± 0.5 and 11.4 ± 0.6, respectively (n= 149). The current traces in Fig.5 A, B, C also indicate that the potassium conductance shows little inactivation in the DCs of CD-1 mice, which contrasts with the pronounced inactivation observed in the DCs of CD-1Cx30^A88V/A88V^ and CBA/J mice. The voltage dependence of activation was shifted to more negative potentials for potassium currents in CD-1Cx30^A88V/A88V^ and CBA/J compared to that of CD-1 DCs (Fig. 5 F). The half-activation voltages (V_1/2_) occurred at significantly (t = 6.81, P < 0.0001, d.f.= 48, α’=0.017) lower potentials in CD-1Cx30^A88V/A88V^ (V_1/2_ =12.4 ± 2.8 mV) DCs than in CBA/J (V_1/2_ =16.5 ± 1.1 mV) DCs. The half-activation voltage in the DCs of CD-1Cx30^A88V/A88V^ was also significantly smaller (t = 22.36, P < 0.0001, d.f.= 48, α’=0.017)) than the voltage measured in the DCs of CD-1 mice (V_1/2_ = 27.9 ±2.3 mV). The maximum slope factor was 18.6 ± 1.2 for CD-1, 21.3 ± 2.0 for CD-1Cx30^A88V/A88V^ and 15.3 ± 0.9 for CBA/J cells. Each point (mean ± SD) is based on measurements for n=20 cells.

**Figure 5.**
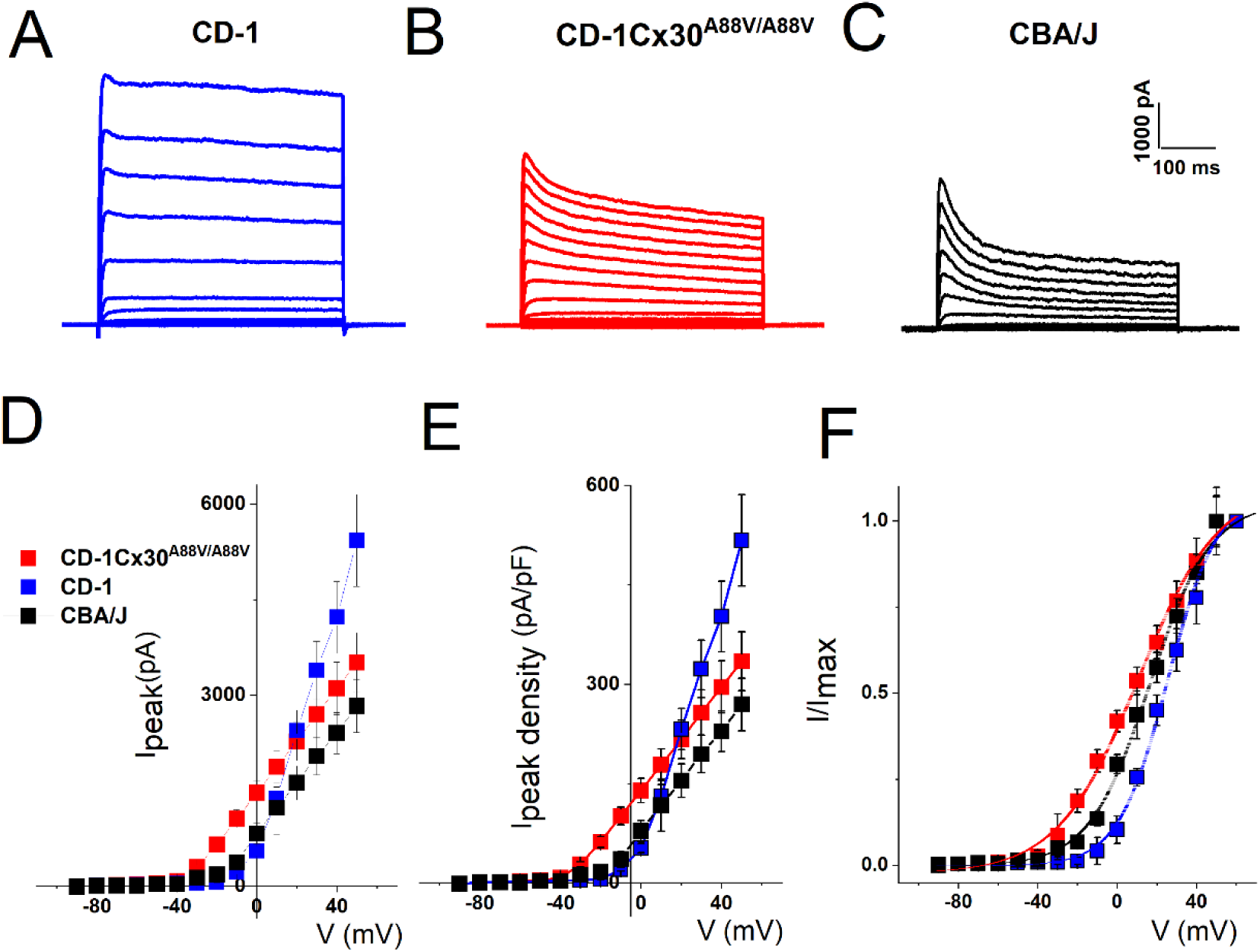
Potassium current amplitude, density, and voltage sensitive activation of CD-1 Deiters’ cells are notably different from to those of CBA/J and CD-1Cx30^A88V/A88V^ Deiters’ cells. Current activation was elicited using 500 ms voltage steps in 10-mV increments from -90 to +50 mV, from a holding potential of -80 mV. Representative examples of current traces recorded from Deiters’ cells (DCs) in: **(A)** CD-1, **(B)** CD-1Cx30^A88V/A88V^ and (**C**) CBA/J mice. The current-voltage **(D)** and current density-voltage **(E)** relationships are notably different between CBA/J, CD-1 and CD-1Cx30^A88V/A88V^ DCs. **(F)** The voltage dependence of activation was assessed for peak potassium currents elicited in CBA/J, CD-1 and CD-1Cx30^A88V/A88V^ DCs. The Boltzman functions were fitted as solid lines.

Together, our results (Fig. 5) show distinct differences in DC potassium currents between CD-1 mice, that exhibit early onset hearing loss, and CD-1Cx30^A88V/A88V^ and CBA/J mice, which have excellent high-frequency hearing.

In order to investigate the possibility that potassium currents in DCs may differ across the different strains, the time- and voltage-dependence of current activation, deactivation and inactivation were examined (Figs. 6, 7 and 8). The current activation was best fitted with a single time constant (in the order of ms, Fig. 6B), but the time constant of CD-1 DCs was about twice those of CBA/J and CD-1Cx30^A88V/A88V^ DCs. There were, however, two different time constants of deactivation (in the order of 10s ms and 100s ms, Fig. 6D) observed for all three mouse strains, indicating the possible presence of multiple currents. This was further reinforced with the studies of the current inactivation behaviour. Voltage-dependence of steady-state inactivation of the transient current was similar in CBA/J and CD-1Cx30^A88V/A88V^ DCs, and steeper than in CD-1 DCs, again, suggesting the possibility of multiple potassium currents (Fig 7). When voltage-dependences of steady-state inactivation were fitted with the Boltzmann function (Fig. 7C), the half inactivation voltages (V_1/2_) (mean ± SD, n=10) were more negative for CD-1 cells (−35.6 ±6.2 mV) than for CD-1Cx30^A88V/A88V^ (−25.9 ± 1.2 mV) and CBA/J (−26.5 ± 1.4 mV) cells. The maximum slope factor (mean ± SD, n=10) for CD-1 and CD-1Cx30^A88V/A88V^ cells were similar, being 6.2 ± 0.6. and 5.9 ± 0.6, respectively. The maximum slope factor was 3.7 ± 0.1 for CBA/J cells. Time constants of inactivation (Fig. 8A, B) and the current recovery from inactivation (Fig. 8C, D) were best fitted with 2 time constants for CD-1, and 3 time constants for CD-1Cx30^A88V/A88V^ and CBA/J DCs. For example, the time constants of inactivation for the depolarizing pulse of +30 mV were (in ms, mean ± SD, n = 12) 170 ± 43 and 7400 ± 360 for CD-1; 65 ± 15, 820 ± 70 and 6387 ± 286 for CD-1Cx30^A88V/A88V^, and 55 ± 10, 620 ± 70 and 7200 ± 430 for CBA/J DCs (Fig. 8B). Also, the time constants for recovery from inactivation (Fig. 8D) were (in ms, mean ± SD, n = 12)) 395 ± 62, 1445 ± 852 for CD-1; 17 ± 2, 175 ± 25; 2965 ± 125 for CD-1Cx30^A88V/A88V^, and 16 ± 2, 180 ± 25; 3400 ± 200 for CBA/J DCs. However, time constants for the development of inactivation were best fit with two time constants in all strains (Fig. 8F). For example, the time constants for depolarizing pulse of +30 mV were (in ms, mean ± SD, n = 12) 340 ± 67 and 6200 ± 260 for CD-1, 330 ± 44 and 6807 ± 256 for CD-1Cx30^A88V/A88V^, and 330 ± 40 and 6400 ± 340 for CBA/J DCs. Therefore, our results suggest that potassium current expression differed between CD-1 DCs, and CD-1Cx30^A88V/A88V^ and CBA/J DCs.

**Figure 6.**
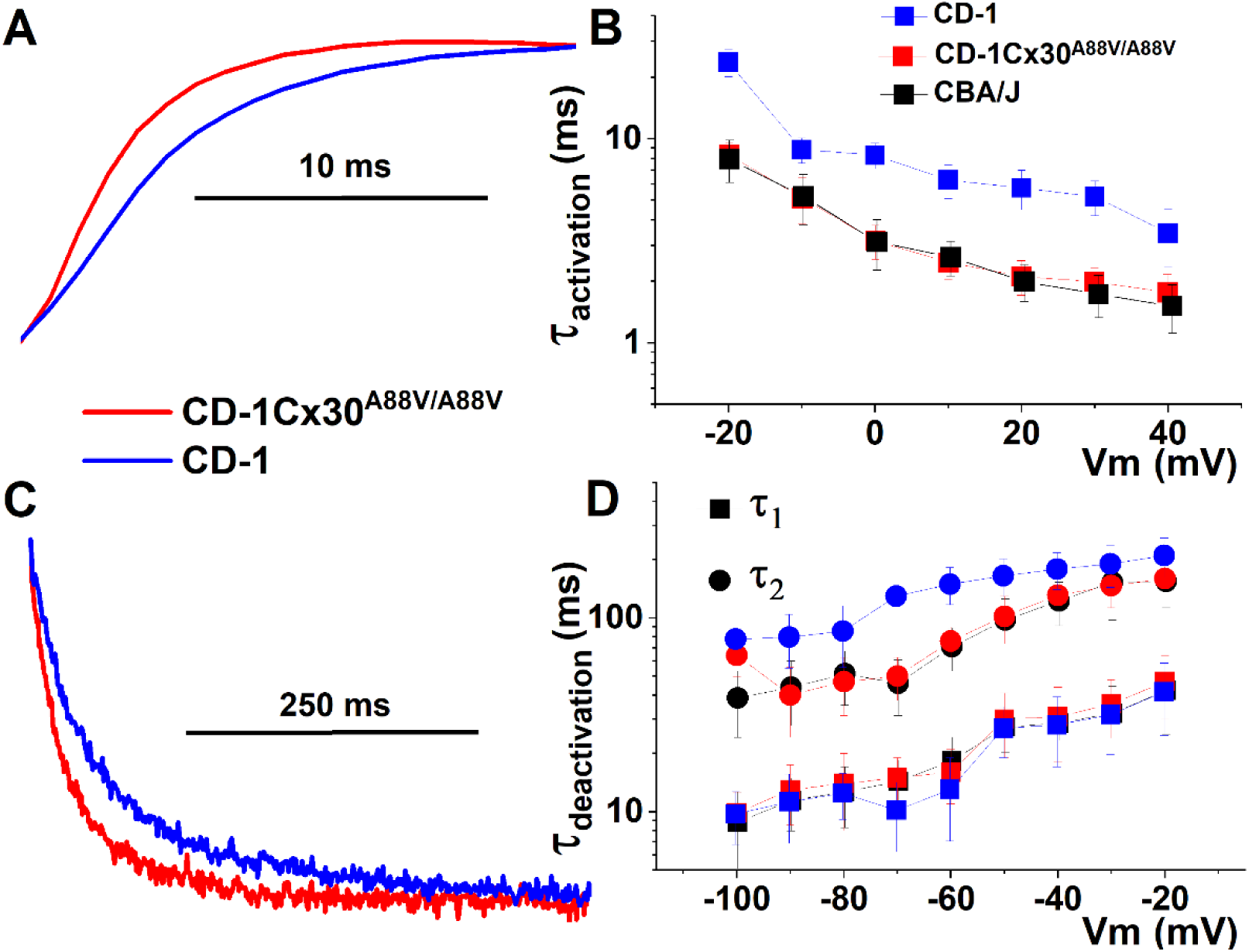
The voltage-dependent time constants of activation of potassium currents are similar in CBA/J and CD-1Cx30^A88V/A88V^ Deiters’ cells, and they are faster than in CD-1 Deiters’ cells and reveal the possibility for multiple potassium conductances. **(A)** Activation of current was elicited with the protocol as in Figure 5. Examples of normalized current traces in CD-1 and CD-1Cx30^A88V/A88V^ Deiters’ cells (DCs) elicited to +30 mV from holding potential of -80 mV. **(B)** The activation time constant was on the order of miliseconds in all three strains but the time constant of CD-1 DCs was about twice those of CBA/J and CD-1Cx30^A88V/A88V^ DCs. **(C)** Tail currents were elicited using 15 ms activating voltage step to +30 mV from a holding potential of -80 mV, followed by a 300-ms test pulse, from -120 mV to -20 mV in 10-mV increments, during which the deactivation time constant was measured. Examples of normalized tail current traces in CD-1 and CD-1Cx30^A88V/A88V^ DCs **(D)** Deactivation time constants of K+-currents for CD-1, CD-1Cx30^A88V/A88V^ and CBA/J DCs. There were two different time constants of deactivation. One in the 10 of ms range, and the other about 10 times longer, for all three mouse strains, indicating the possible presence of multiple currents.

**Figure 7.**
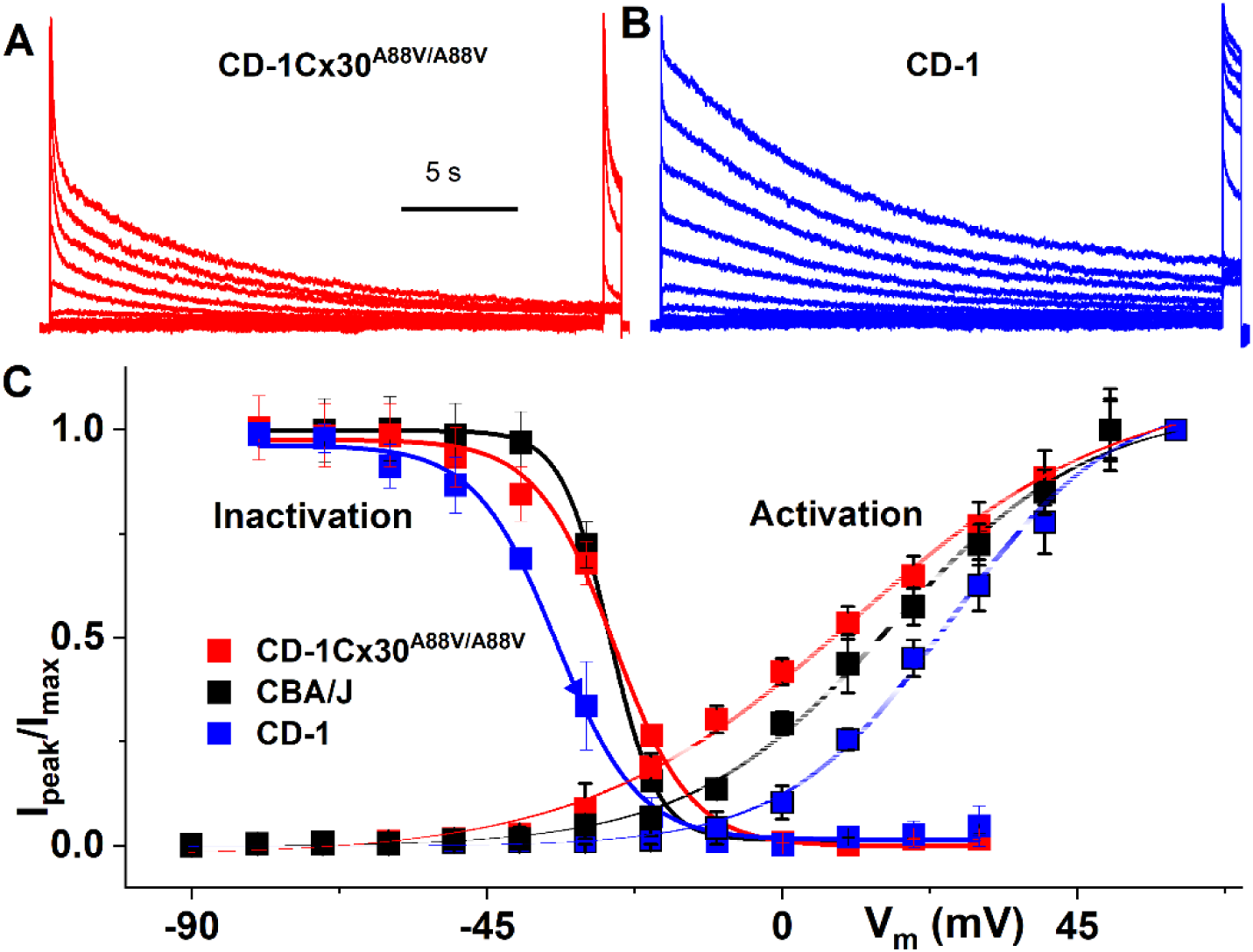
Voltage-dependence of potassium current steady-state inactivation for Deiters’ cells is similar in CD-Cx30^A88V/A88V^ and CBA/J and steeper than in the CD-1 cochleae. Steady-state inactivation properties of the transient current were determined by presenting pre-pulses of 16-sec duration, at different membrane potentials (−90 to +30 mV), followed by a test pulse at +30 mV for 300 ms. Examples of current traces recorded from CD-1Cx30^A88V/A88V^ **(A)** and CD-1 **(B)** Deiters’ cells. **(C)** Normalized dependence of I_peak_ during the test pulse showing voltage dependent inactivation. I_max_ is I_peak_ for -90-mV pre-pulse. The Boltzmann function fits for steady-state inactivation are plotted with solid lines. Each point (mean ± SD) is based on measurements for n = 10 cells. Panel (C) also shows voltage-dependence of activation replotted from Figure 5F. The window current became evident from the intersection of activation and inactivation curves, suggesting that the potassium currents may be partially activated at rest.

**Figure 8.**
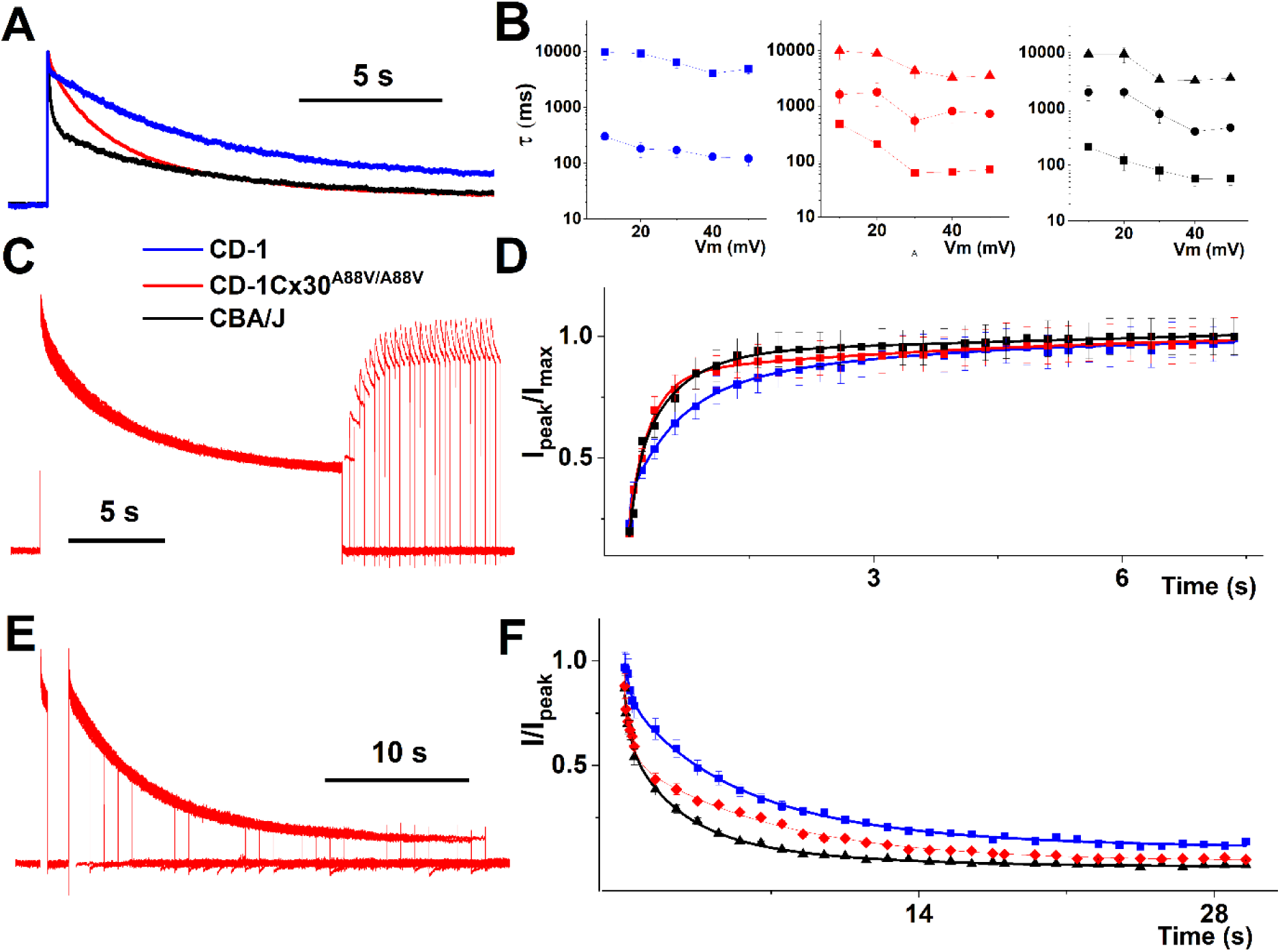
Time-dependence of potassium current inactivation in Deiters’ cells suggests two different potassium channels in CD-1 and three different potassium channels in CBA/J and CD-1Cx30^A88V/A88V^ cochleae. **(A)** Example of normalized current traces elicited by 15 sec depolarizing pulses from holding potential of -80 mV to +30 mV illustrate different time-dependence of inactivation for CD-1, CD-1Cx30^A88V/A88V^ and CBA/J. **(B)** When potassium currents were elicited by 15 sec depolarizing pulses from holding potential of -80 mV to 0 – 50 mV in 10 mV increment, the current inactivation could be best fitted with 2 time constants for CD-1 and with 3 time constants for CD-1Cx30^A88V/A88V^ and CBA/J Deiters’ cells (DCs). **(C)** The potassium current recovery from inactivation was assessed using a standard recovery protocol. Namely, inactivation was caused by a 15-second depolarizing pulse from the -80 mV holding potential to +30 mV. At the end of this conditioning pulse, the cell’s potential was returned to the holding potential of -80 mV for a variable time. The recovery from inactivation was estimated measuring the potassium current amplitude elicited by a test depolarising pulse to +30 mV at the end of the variable time at the holding potential of -80 mV and plotting it against the variable duration of steps to -80 mV. **(D)** Normalized peak potassium current during the test pulse of +30 mV as a function of the variable time of the holding potential at -80 mV preceding the pulse (protocol (C)). This dependence shows the inactivation recovery kinetics and is best fitted with 2 time constants for CD-1 and with 3 time constants for CD-1Cx30^A88V/A88V^ and CBA/J DCs. **(E)** Protocol to examine the development of inactivation kinetics. Depolarizing steps of variable duration from the holding potential of -80 mV to +30 mV were used to elicit potassium currents. The currents were measured at the end of each pulse. **(F)** Normalized potassium currents measured using protocol (E) plotted against the duration of the step to +30mV. Each point in B, D, F, represents mean ± SD for n=12 cells.

Thus, these findings suggest that inactivating potassium current is more dominant in CD-1Cx30^A88V/A88V^ and CBA/J DCs, both with excellent high frequency hearing, whereas the sustained component prevails in CD-1 mice with early onset hearing loss.

## DISCUSSION

Measurements presented here reveal that the A88V/A88V mutation of the GJ protein Cx30 in the CD-1 mouse strain causes electrophysiological changes in the DCs and interconnecting GJs that rescues the phenotypical, early-onset presbycusis of the CD-1 mouse strain (Figures 1-4). This is despite causing reductions in EP and low-frequency hearing sensitivity compared with the wild-type CD-1 mouse. Significantly, the mutation also alters the expression and properties of the DC potassium conductances so that they closely resemble those of CBA/J mice, with excellent high-frequency hearing, rather than those of the CD-1 background strain (Figures 5-8). Both outcomes provide a basis for understanding the ability of mice, and perhaps other mammals, to hear high-frequency sounds that can exceed frequencies of 100 kHz.

### Increased organ of Corti resistance in CD-1Cx30^A88V/A88V^mice, compared with CD-1 and CBA/J strains, is associated with increased input resistance and weaker syncytial junctional coupling in Deiters’ cells

From previous *in vivo* measurements, the rescue of high-frequency hearing loss in CD-1Cx30^A88V/A88V^ mice was proposed to be due to a measured increase in the resistance of the OoC, which was attributed to the Cx30 A88V/A88V mutation expressed in DCs and outer pillar cells (Levic et al., 2022). The OHC ERPs, which are not attenuated at high frequencies, dominate transmembrane potentials across the OHC basolateral membranes and, thus drive prestin-mediated OHC electromotility at high-frequencies (Levic et al., 2022). The higher OoC resistance results in the OHC ERPs being of similar size in CD-1Cx30^A88V/A88V^ and CBA/J mice despite smaller EP and, hence, smaller MET current across the OoC resistance. Findings reported here of increased input resistance and weaker syncytial junctional coupling in DCs cells from CD-1Cx30^A88V/A88V^ mice (Fig 1) support this proposal. Moreover, heterogeneous syncytial junctional coupling was absent in these mutants (Zhao, 2000; Zhao & Santos-Sacchi, 2000). The A88V/A88V mutation of Cx30 does not appear to be able to form heterotypic GJs with Cx26, an inability that has consequences for the generation of EP. This might be expected since the mutated Cx30A88V is incapable of forming functional GJs (Berger et al., 2014). Thus, syncytial junctional conductance is determined only by homotypic GJ coupling in DCs from CD-1Cx30^A88V/A88V^ mice (Fig. 4), and the size of supporting cell Cx30 GJs in CD-1Cx30^A88V/A88V^ mice and overall expression of Cx30 GJs is reduced in the cochlea, including in regions associated with the generation of the EP (e.g. fibrocytes, (Kelly et al., 2019)). Positive EP is generated in the cochlear lateral wall by marginal cells and the intermediate cells coupled with the basal cells in the stria vascularis and neighbouring fibrocytes in the spiral ligament by GJs (Nin et al., 2008). Loss of heterogeneous GJs in these epithelial cells is suggested to cause reduced EP (Mei et al., 2017). This is also observed in Cx30^A88V/A88V^ mice (Levic et al., 2022) and associated with their poor low-frequency hearing, which appears to be driven by intracellular RPs which are reduced as a consequence of the reduced EP.

Heteromerization of GJ proteins is believed to enable the fine regulation of cell-to-cell communication (Cottrell & Burt, 2001) but is absent in Cx30^A88V/A88V^ mice ((Berger et al., 2014), see above). This absence does not appear to adversely influence high-frequency hearing or protection against presbycusis (Kelly et al., 2019). Findings reported here for *ex vivo* measurements from DCs also accord with those from *in vitro* cultured cells and cell pairs in that the expression of Cx30A88V, rather than Cx30, causes a reduction in the conductance of GJs and changes their voltage gating properties (Berger et al., 2014; Essenfelder et al., 2004).

### Deiters’ cell potassium conductances in CD-1Cx30^A88V/A88V^mutants have closely similar properties to those of CBA/J mice with excellent hearing and differ from those of wild-type CD-1 mice

Whole-cell patch recording from DCs in isolated sections of the OoC reveal similarities in the K^+^ conductances of DCs from CD-1Cx30^A88V/A88V^ and CBA/J mice, both with excellent high-frequency hearing and without early-onset presbycusis. The DC K^+^ conductances differ, however, from those measured in the wild-type CD-1 mice with poor high-frequency hearing and early-onset presbycusis. Our finding indicates the existence of several K^+^ channel types that can be expressed in different ratios and perhaps distributions within the DCs of the different mouse strains. Differences in K^+^ channel types and distribution could be due to differences in ionic and electrical environments during development caused by expression of different GJs and their patterning (Sousonis et al., 2021).

The K^+^ conductances are activated when the DCs are depolarized. *In vivo*, this is presumably during sound stimulation when K^+^ levels in the surrounding Cortilymph are increased and this extracellular fluid space is depolarised (Cody & Russell, 1987; Johnstone et al., 1989). The half-activation voltages of DC K^+^ conductances in CD-1 mice are more positive than in CD-1Cx30^A88V/A88V^ and CBA/J mice. However, the threshold for activation in CD-1Cx30^A88V/A88V^ is more negative than in CBA/J and CD-1 mice. These observations could indicate that CD-1Cx30^A88V/A88V^ mice are most sensitive to increases in the positivity of potentials in the Cortilymph, while CD-1 mice are relatively insensitive to these changes. The DC K^+^ conductances are also slower to activate and inactivate, and the K+ current density and amplitude are larger in CD-1 than those in CD-1Cx30^A88V/A88V^ and CBA/J mice. Moreover, the K^+^ conductance poorly inactivates in wild-type CD-1 mice. It may be that the relative insensitivity of the DC K^+^ conductance of CD-1 mice to increases in positivity of the Cortilymph and their relatively slow responses to changes in membrane potential have consequences for the cochlear K^+^ recycling and early-onset presbycusis (Spitzmaul et al., 2023). The rapid, more sensitive responses of the DC K^+^ conductance in CD-1Cx30^A88V/A88V^ mice, which resemble responses in CBA/J strain, may contribute to the sensitive, high-frequency hearing in these mice and their resistance to presbycusis.

It is widely accepted that the resting potential of specific cells is set by the properties of ionic conductances, and GJs regulate the topology of tissue-level bioelectric networks by regulating the boundaries between cell fields with distinct transmembrane potential states (Levin, 2021). Findings reported here, and previously (Levic et al., 2022), do not entirely support this concept. The resting potentials of presumed DCs in CD-1Cx30^A88V/A88V^ and CBA/J mice, are not significantly different (Levic et al., 2022) being close to -110 mV. This is close to the activation voltage of the K^+^ conductance. Thus, the resting membrane potentials of DCs in CD-1Cx30^A88V/A88V^ and CBA/J mice appear not to be dominated by the electrophysiological properties and density of the GJs and potassium conductances, and maybe controlled, for example, by K/Cl cotransporters (Boettger et al., 2002; Boettger et al., 2003; Zdebik et al., 2009). The resting membrane potentials of supporting cells measured *in vivo*, at least from mice and guinea pigs (Russell & Sellick, 1978), for example, are more negative than the OHCs they encompass. Unfortunately, it was not possible to measure DC resting potentials, *in vivo*, from the high-frequency region of the CD-1 mouse, due to degeneration of this region associated with high-frequency hearing loss.

### Significance of electrophysiological properties and distribution of organ of Corti gap junctions for cochlear mechanical and electrical interaction

It might be expected that reduction in the conductance, size, and distribution of Cx30 GJs due to mutation in CD-1Cx30^A88V/A88V^ mice should also cause changes in the extent to which electrical, ionic, and even molecular, changes in one part of the cochlea can influence voltage and consequently mechanical responses in adjacent regions. A test of this would be to measure phenomena, including two-tone suppression (Delgutte, 1990; Ruggero et al., 1992; Sellick & Russell, 1980) and the longitudinal spread of excitation along the reticular lamina (Ren et al., 2016), which could be different in CD-1Cx30^A88V/A88V^ mice compared with, say, that of CBA/J mice, although cochlear sensitivity to high-frequency tones of these two mouse strains is similar. Such a test would also reveal the extents to which the local cochlear responses are determined directly by the local mechanical properties of the cochlea and indirectly through the electrical interaction (e.g. (Mistrík et al., 2008)) and consequences of electrical changes in the supporting cells and interstitial spaces of the OoC.

### Cx30A88V expression and presbycusis

The mechanisms by which the expression of Cx30A88V staves-off presbycusis remains a mystery. (Kelly et al., 2019) concluded Cx30A88V expression could preserve high-frequency hearing in old age and preservation of high-frequency hearing was associated with a reduction in Cx30, but not Cx26. Measurements presented here provide additional possibilities including the inability of Cx30A88V to combine with Cx26 to provide heterozygous GJs and reduced connectivity between supporting cells. This is significant if, as has been shown for noise induced hearing loss (Xiao et al., 2025), supporting cells, including DCs, orchestrate hearing loss through a feedback pathway whereby noise-induced oxidative stress in OHCs initiates an OHC to supporting cell cascade involving Gasdermin D (GSDMD). When activated in supporting cells, GSDMD exacerbates oxidative injury in both cell types via a reciprocal pathogenic positive feedback loop. The loop might be impaired through Cx30A88V expression, and reduced intercellular communication between supporting cells, thereby preserving high-frequency hearing. Another possibility involves the poor low-frequency hearing of CD-1Cx30^A88V/A88V^ mice. Exposure to low-frequency noise in their environment (Lauer et al., 2009; Reynolds et al., 2010) has been shown to exacerbate the occurrence of presbycusis in rodents (Liu et al., 2022; Sun et al., 2025). Perhaps expression of Cx30A88V, and poor low-frequency hearing, contributes to preserving high-frequency hearing in elderly Cx30A88V mice.

## ADDITIONAL INFORMATION

### Data availability statement

Raw data will be made available to any qualified scientists on reasonable request.

### Author contributions

All experiments were performed in the Sensory Neuroscience laboratory at the University of Brighton, Brighton, UK. A.N.L., S.L., I.J.R. conceived and coordinated the study and wrote the paper. P.S. and S.L. were primarily responsible for data collection and analysis, and all other authors contributed to this. V.A.L. and P.S. edited the final version of the paper. A.N.L. and I.J.R. raised funding. All authors agreed to be accountable for all aspects of the work in ensuring that questions related to the accuracy or integrity of any part of the work are appropriately investigated and resolved. All persons designated as authors qualify for authorship, and all those who qualify for authorship are listed.

## Authorship

S. Levic and I. J. Russell share last authorship.

